# The impact of physiological state and environmental stress on bacterial load estimation methodologies for *Mycobacterium tuberculosis*

**DOI:** 10.1101/2024.05.10.593528

**Authors:** Arundhati Maitra, Marie Wijk, Hasmik Margaryan, Paolo Denti, Timothy D. McHugh, Frank Kloprogge

## Abstract

**Background:** Solid and liquid medium cultures from patient samples recover different proportions of a heterogenous bacterial community over the duration of treatment. *In vitro* experiments were designed to study the population composition at early-logarithmic and stationary phases of growth as well as under drug pressure.

**Objectives:** To derive a relationship between methodologies for bacterial load determination and assess the effect of the growth phase of the parent culture and its exposure to stress on the results.

**Methods:** *Mycobacterium tuberculosis* H37Rv was grown with and without drug (isoniazid or rifampicin) and sampled on day 0, 3, 11 and 21 of growth in broth culture. The bacterial load was estimated by colony counts and the BD BACTEC™ MGIT™ automated mycobacterial detection system. Linear and nonlinear mixed-effects models were used to describe the relationship between time-to-positivity (TTP) and time-to-growth (TTG) vs colony forming units (CFU), and growth units (GU) vs time.

**Results:** For samples with the same CFU, drug-treated and stationary phase cells had a shorter TTP than the drug-free control and early-logarithmic phase cells respectively. Similarly, stationary phase samples reached higher GUs and had shorter time to start growing than early-log phase ones.

**Conclusions:** The growth phase affects the relationship between CFU-TTP/TTG and previous exposure to drugs affects only the relationship between CFU-TTP. This suggests that there is a population of bacterial cells that can be differentially recovered in liquid medium giving us an insight into the physiological states of the original culture which aids in the interpretation of clinical trial outputs.

## Introduction

As tuberculosis (TB) remains a global health crisis, infecting over 10 million individuals and claiming the lives of around 1.6 million patients annually, methodologies to identify efficacious drugs and drug combinations remains a priority.^1^

Since the landmark publication by Jindani 1980, early bactericidal activity (EBA), or the reduction of bacterial load in patient samples over the first two weeks of treatment, has been the cornerstone of estimating anti-mycobacterial efficacy of novel drugs and drug combinations.^2,3^ Historically, colony counts were the sole means to estimate bacterial loads. Colony forming units (CFU) are estimated based on direct observation of bacterial growth. This relies upon the ability of the bacteria to multiply on solid agar medium. Time-to-positivity (TTP), on the other hand, is an indirect measure of bacterial load based on oxygen consumption in the Mycobacteria Growth Indicator Tube (MGIT). The rate and/or amount of oxygen consumption is a function of the bacterial load as well as the physiological state of the bacilli namely, actively replicating versus metabolically dormant. TTP is defined as the incubation time required to reach a threshold set by the manufacturer and measured in growth units (GU). Several groups have validated the use of TTP as a robust replacement for the CFU measurement, considering its reproducibility and ease-of-use.^4–6^ Additionally, the REMoxTB trial demonstrated that MGIT culture can replace solid culture in phase III trials but did note that there may be implications for the primary endpoint analysis.^7^

Studies have shown that *M. tuberculosis* exists as a heterogeneous population and recovery of the bacilli depends on the type of media. Liquid media allows for the growth of a population subset that does not grow on solid media owing to phenotypic differences in the pathogen.^5,8^ Supplementation of liquid media with resuscitation-promoting factors (RPFs) has been shown to result in even higher recovery rates highlighting the presence of several phenotypic states of the bacteria and the importance of determining the relative proportions of these populations during treatment and how their elimination correlates with treatment success or the relapse of infection.^9^

Diacon *et al*. and Bark *et al*. studied the relationship between CFU and TTP in samples collected from 250 and 41 patients respectively over the first 14 days of treatment.^5,6^ Both studies identified a negative correlation that was linear at lower CFU counts (below 5 logCFU).

Bowness *et al*. followed 68 patients and in addition to the negative correlation between TTP and CFU, found that as treatment progressed, samples with identical CFU had longer TTP.^10^ This could indicate that as treatment progresses, (a) the proportion of bacteria additionally picked up by the TTP measure (ie., recovered in liquid media) is reduced or (b) for the same number of cells, the treated cells had lower metabolic activity. The patients in the study received rifampicin monotherapy at different doses for the first 7 days followed by standard therapy with isoniazid, pyrazinamide and ethambutol. A dose-dependent effect was observed indicating that rifampicin was effective in altering the TTP readings. However, an orthogonal means of measurement is required to assess whether this is due to changes in the bacterial load or the basic physiology of the pathogenic population recovered.

It has been observed that the linearity of the relationship between TTP and CFU breaks down at high bacterial loads, over the course of treatment, as well as with increasing doses of antimicrobials. Additionally, in the patient sample studies described above, the effect of the drug on the heterogenous populations of the pathogen cannot be unravelled from the effect of its physiological growth phase. A mechanistic understanding of the types of populations found and the dynamics between them at each phase of growth and under drug pressure is required. Therefore, we designed *in vitro* experiments to study the effect of the incubation period, or the growth phase of the culture in the presence or absence of drugs so that for every drug-containing sample we had a matched drug-free control. This would enable us to identify the changes in the population dynamics, i.e., populations that are differentially detected in liquid culture, as a function of the length of the incubation period and identify how the addition of drug pressure affects it.

## Materials and Methods

### In vitro experiments

*Mycobacterium tuberculosis* H37Rv (ATCC 25618) was used for all the experiments. The seed culture was grown for 6 days (early-log phase) in BD BACTEC™ MGIT™ (Cat No. 245122) containing SIRE supplement (800 μL of SIRE supplement in 7 mL tubes). All reagents were procured from Merck unless otherwise mentioned.

10 mL Middlebrook 7H9 broth containing 0.02% (*v/v*) glycerol, 0.05% (*v/v*) Tween-80 supplemented with 10% SIRE supplement in a glass universal bottle was inoculated with 0.1 mL of the early-log phase culture and incubated as a stand culture at 37 °C. On days 3, 11 and 21 post inoculation, samples were withdrawn in duplicate and diluted ten-fold. From each dilution, 100 μl was used to spread on a Middlebroook 7H10 plate (for CFU counts) and 100 μL was used for inoculation into a MGIT tube (for TTP determination). There were three biological replicates, each with two technical replicates within the experiment.

For the drug containing experiments, 1/2x MIC of isoniazid and rifampicin (0.1 μg/mL and 0.0015 μg/mL respectively) was added on the day of the inoculation and sampled as mentioned. There was one experiment performed per drug and one experiment without drugs, all with two technical replicates.

Plates were checked for growth after 14, 21, 28 days and the CFU was recorded as the number of colonies counted. GU is the output measure provided by BD BACTEC™ MGIT™ systems as a means of bacterial load estimation. It is derived from the fluorescence emitted by the embedded oxygen-quenched fluorochrome in the tubes. As the dissolved oxygen reduces due to metabolic activity or active replication of cells the fluorescence signal, and in turn GU, increases. TTP is the time required for a MGIT to reach a predetermined GU as set by the manufacturer. Both TTP and hourly GU measurements were recorded by the BD EpiCenter™ Microbiology Data Management System.

#### Analysis of relationship between TTP and CFU

Linear mixed-effects modelling in NONMEM version 7.5.0 was used to describe the relationship between TTP and CFU.^11^ Pirana, Pearl-speaks-NONMEM, and R version 4.2.0 were used to assist the modelling process.^12–14^ TTP in hours on natural logarithmic scale was treated as the dependent variable and CFU on log10 scale as the independent variable. An additional dataset was created in which all CFU data, including missing values, were imputed based on the geometric mean of the dilutions of the same sample with readable CFUs. Linear models estimating an intercept and a slope were developed, with a variability component separating random variability between different biological replicates from residual variability. The intercept was centered around 1 log(10) CFU, at which there was more data available, rendering the estimation more stable. Random variability was tested on both intercept and slope, assumed to be additive on natural logarithmic scale for the intercept and log-normally distributed for the slope. An additive error model on logarithmic scale was tested to describe the residual variability. Error may be introduced while reading crowded plates with merged colonies, i.e. >100 CFU. Similarly, samples with CFU <10 are prone to error due to the clumping tendency of mycobacterial cells. Therefore, additional error was tested for samples with a CFU <10 or >100. Covariates were tested using a stepwise approach with forward inclusion (p ≤ 0.05) followed by backwards elimination (p ≤ 0.01). The inclusion of covariates was guided by objective function value (OFV), visual predictive checks (VPC) and goodness-of-fit plots. Covariates tested were growth phase of the culture and the presence of drug.

#### Analysis of relationship between time-to-growth and CFU

The relationship between time-to-growth (TTG), defined as the time taken to obtain the first GU reading greater than 0, and CFU was analyzed with linear mixed-effects modelling, using same software as for the TTP and CFU analysis and CFU values were imputed in an additional dataset in the same manner. TTG in hours on natural logarithmic scale was treated as the dependent variable with CFU on log10 scale being the independent variable. A linear model estimating an intercept and a slope was developed to describe the relationship between TTG and CFU with and without imputations. The intercept was centered around log(10) CFU, and random variability, residual variability and covariates were included in the same manner as for the TTP and CFU analysis.

#### Analysis of relationship between GU and incubation time in MGIT

The relationship between GU and incubation time in the MGIT was analyzed with nonlinear mixed-effects modelling, using the software mentioned. The natural logarithm of GU was used as the dependent variable and incubation time in the MGIT was used as the independent variable. The TTG was subtracted from each sequence of GUs to compare the growth trajectories after growth detection only. Exponential, logistic, and E_max_ functions were tested to fit the data (equations below).

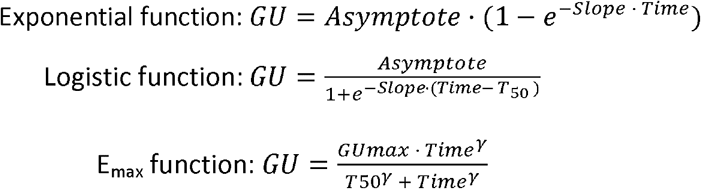

The asymptote in the exponential and logistic function, and GU_max_ in the E_max_ function represent the carrying capacity of the growth, i.e. where the growth curve reaches a plateau. The slope in the exponential and logistic function represents how fast the bacteria grow. The parameter γ in the E_max_ function determines the shame of the GU curve over time. T50 is the time at which half of the carrying capacity is reached.

A variability component separating random variability between GUs from each MGIT from residual variability was introduced. Random variability was assumed to be either additive on natural logarithmic scale or log-normally distributed and tested on all estimated parameters, and covariance between random variabilities was estimated where correlation was indicated in diagnostic plots. An additive error model on logarithmic scale was tested to describe the residual variability. Growth phase of the culture and presence of drug were tested as covariates in the same manner as for the models for TTP or TTG and CFU.

## Results and Discussion

### Analysis of relationship between TTP and CFU

In total, 110 matching TTP and CFU samples and an additional 84 imputed CFU samples with matching TTP were included in the analysis. As expected, a negative relationship between TTP and CFU was observed, which was adequately described by a linear model estimating an intercept and slope in both datasets (Figure 1 and S1, Table 1 and S1, and Suppl. A). Results from the data with imputations on indeterminate CFUs can be found in supplementary material. In the dataset without imputations, all samples other than those taken at day 3 had 18.6-folds larger random variability. This suggests that the inoculum taken from a heterogenous population undergoes adaptive processes in the initial phase of growth (around day 3) that results in the lowering of the physiological heterogeneity. Following this, due to stochastic processes such as uneven cell divisions, the population reverts to being heterogenous.^15–21^

**Figure 1.**
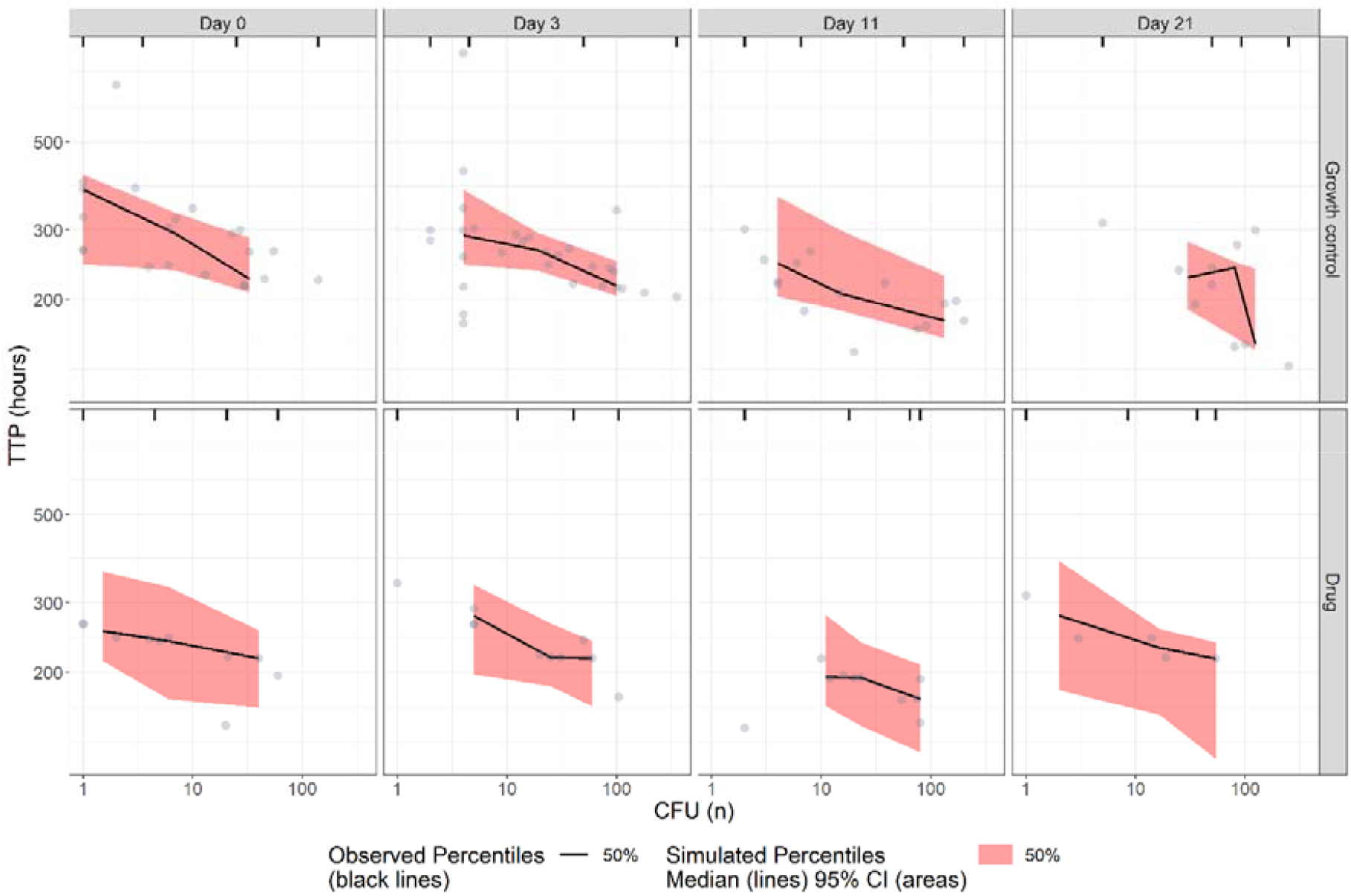
Visual predictive check (n=1000) of TTP versus CFU without imputations, stratified by culture-age and presence of drug. The circles represent observations.

**Table 1:** Parameter estimates for model describing relationship between TTP and CFU without imputations.

| Parameter | Typical value (95% CI)<sup>a</sup> |
| --- | --- |
| Intercept (hours) | 215 (198 – 232) |
| Slope (hours/log10 CFU) | -50.3 (-65.5 – -35.2) |
| Additive error on log scale (%) | 13.7 (11.7 – 16.9) |
| Scaling of additive error on log scale for < 10 CFU (folds) | 1.94 (1.48 – 2.51) |
| Between-biological replicate variability on intercept (%CV) | 0.48 (0.23 – 0.66) |
| Scaling of random variability on intercept for culture-age 0, 11 or 21 days (folds) | 18.6 (8.00 – 31.9) |
| Effect of drug presence on intercept (%) | -17.6 (-27.3 – -7.73) |
| Effect of culture-age 11 or 21 days on intercept (%) | -15.3 (-22.8 – -7.21) |

<sup>a</sup> 95% confidence interval obtained by sampling importance resampling (SIR) using Pearl- Speaks-NONMEM.

Additive error was inflated by 1.94-fold for samples with <10 CFUs reflecting that clumping of cells can significantly impact the CFU counts, especially in samples with low population densities. Accounting for the additional error introduced when reading plates with >100 CFU was not as statistically significant as inflating the error for samples with <10 alone, nor did it improve the diagnostic plots.

For samples with the same CFU, drug-treated cultures had a shorter TTP than control. Similarly, samples with the same CFUs withdrawn from stationary phase cultures had shorter TTP compared to samples withdrawn from early-logarithmic phase cultures. This indicates that with longer incubation periods or in the presence of drugs, either the proportions of bacterial cells that can be recovered on solid medium decrease or the metabolic activity of the population is altered. In the first scenario it would mean that the populations recovered in solid and liquid media in the untreated or early-logarithmic phase of growth largely overlap. Whereas, as the culture experiences drug pressure or progresses to stationary phase, bacterial populations arise that preferentially grow in liquid over solid media. Without a measure of the metabolic activity levels of cells from both the cultures when resuspended in fresh MGIT broth, the impact of the difference in cellular physiology also cannot be ruled out.

### Analysis of relationship between TTG and CFU

TTG represents the first time point at which GU is above 0. Apart from the limit of detection of the instrument, this period also includes the lag phase wherein the cells prepare to divide. We investigated this early stage to identify whether previous drug exposure or physiological growth phase of the parent culture affected recovery in liquid media.

In total, 90 matched TTG and CFU reads were included in the analysis from 5 out of 6 biological replicates, as the whole GU trajectory was not available for all biological replicates. An additional 71 imputed CFU values with matching TTG were included in the dataset with imputations, the results from which can be found in the supplementary material (Figure S2, Table S2, Suppl. B). A linear model adequately described the relationship between TTG and CFU (Figure 2, Table 2, Suppl. B). The physiological growth phase of the culture significantly affected TTG. For the same CFU, TTG was found to be shorter for samples obtained from stationary phase cultures than early-logarithmic phase cultures. In other words, for the same TTG, a higher number of cells is recovered on solid media for early-stationary phase cells compared to stationary phase cultures. This could indicate that stationary phase cultures are largely made up of a population that preferentially grows in liquid medium or metabolically recovers faster on being exposed to fresh MGIT broth. The presence of drugs did not have any impact on the relationship between TTG and CFU.

**Figure 2.**
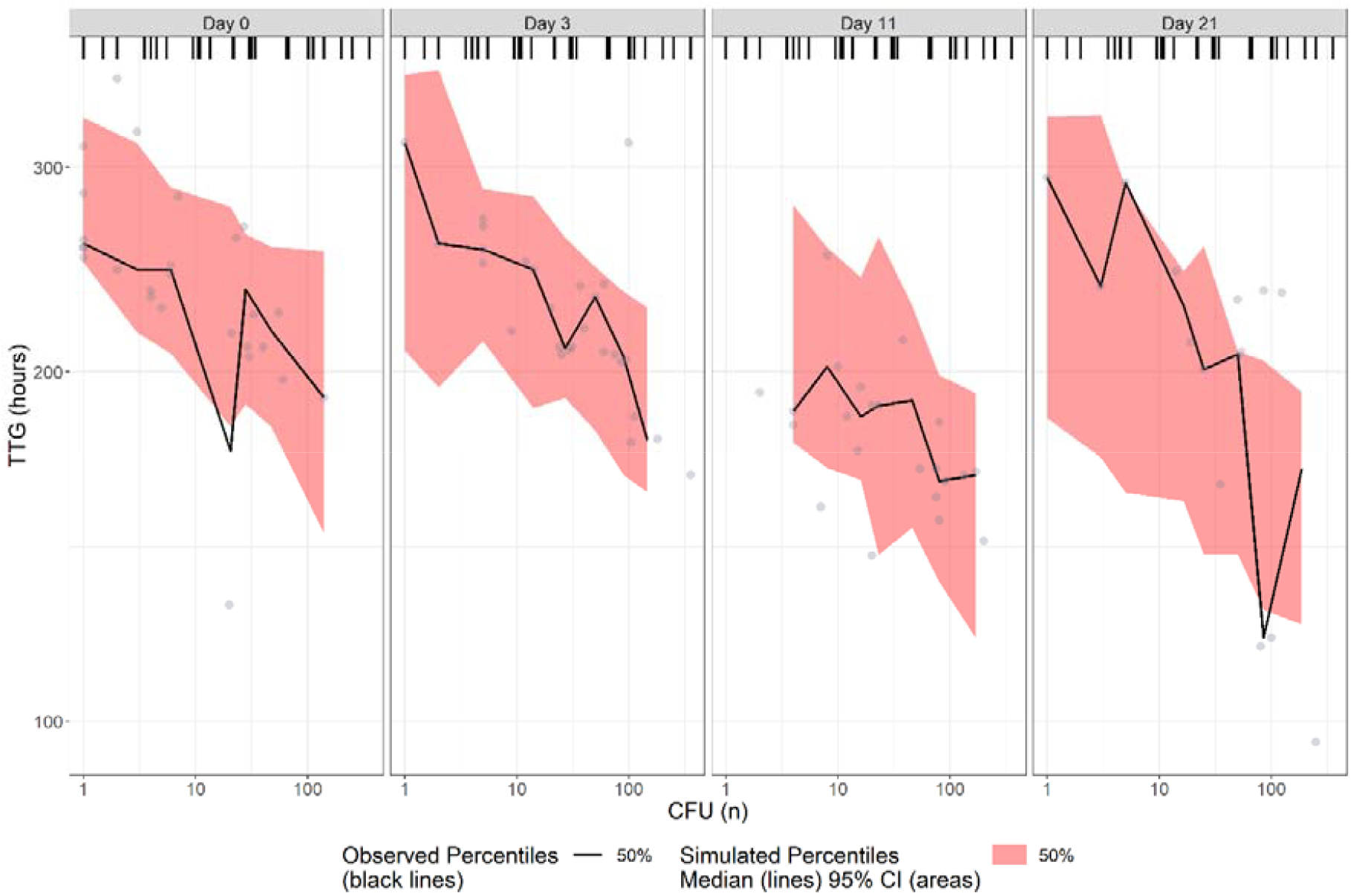
Visual predictive check (n=1000) of TTG versus CFU without imputations, stratified by culture-age. The circles represent observations.

**Table 2:** Parameter estimates for model describing relationship between TTG and CFU without imputations.

| Parameter | Typical value (95% CI)<sup>a</sup> |
| --- | --- |
| Intercept (hours) | 184 (167 – 202) |
| Slope (hours/log10 CFU) | -42.2 (-52.9 – -31.2) |
| Additive error on log scale (%) | 14.6 (12.8 – 17.2) |
| Between-biological replicate variability on intercept (%CV) | 8.33 (4.95 – 11.6) |
| Effect of culture-age 11 and 21 days on the intercept (%) | -18.3 (-24.7 – -11.6) |

<sup>a</sup> 95% confidence interval obtained by sampling importance resampling (SIR) using Pearl-Speaks-NONMEM.

### Analysis of relationship between GU and incubation time in MGIT

A total of 184,129 hourly observations from 227 experiments were included in the analysis (Figure 3, Table 3 and Suppl. C). The relationship between GU and incubation time in MGIT was best described by a logistic model. Growth phase significantly affected the slope and asymptote, with stationary phase samples growing faster and reaching higher GU than the early-logarithmic phase samples. This is not surprising as the samples from later time-points would have higher bacterial loads than those from the earlier samplings. The presence of drug did not affect the relationship between GU and incubation time in MGIT, indicating that once removed from drug pressure, the cells reverted to the same state with similar population kinetics. We attempted to model GU and TTG combined as dependent variable versus time, but reverted to model them separately due to numerical issues.

**Figure 3.**
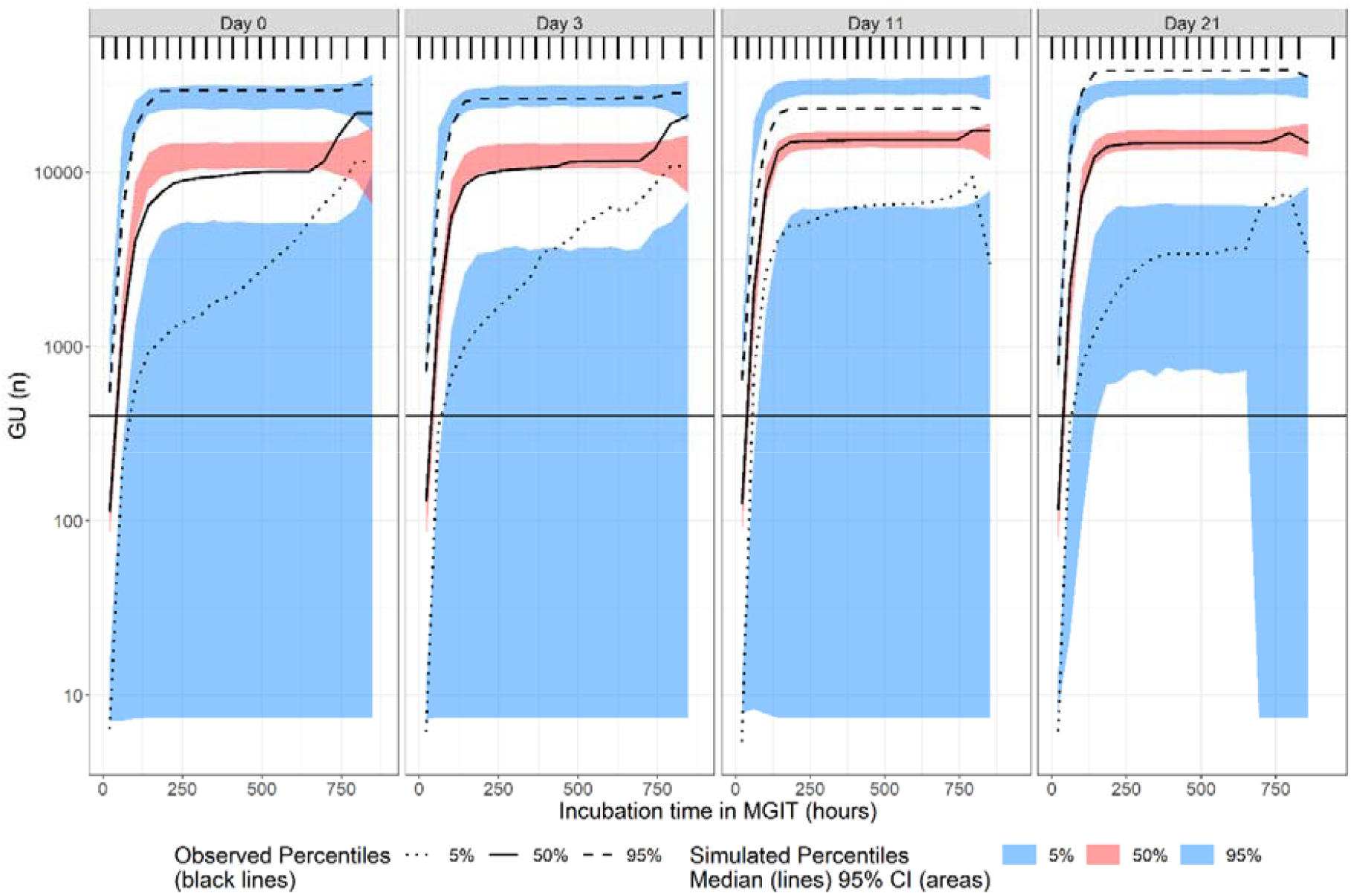
Visual predictive check (n=1000) of GU versus TTP, stratified by culture-age. The vertical line represents the GU around which TTP is usually recorded (GU = 400). The individual observations have been omitted from the plot due to their abundance, rendering the simulations imperceptible. The 5% observed percentiles have wider confidence intervals due to the effect of samples in the higher dilutions. The maximum GU for these samples plateaued at lower levels and these had fewer data points as well, as shown in Figure S3 and Figure S4 in supplementary material.

**Table 3:** Parameter estimates for model describing relationship between GU and incubation time in MGIT.

| Parameter | Typical value (95% CI)<sup>a</sup> |
| --- | --- |
| Asymptote (GU) | 12,800 (11,363 – 14,237) |
| Slope (GU/hour) | 0.069 (0.065 – 0.074) |
| T50 (hours) | 88.9 (84.6 – 93.2) |
| Additive error on log scale (%) | 21.6 (20.3 – 22.9) |
| Between-MGIT variability on asymptote (%CV) | 58.9 (52.3 – 64.6) |
| Between-MGIT variability on slope (%CV) | 48.6 (38.8 – 56.7) |
| Between-MGIT variability on T50 (%CV) | 35.8 (24.7 – 44.1) |
| Correlation between random variability on slope and T50 (%) | -94.8 (-96.5 – -93.1) |
| Effect of culture-age 11 or 21 days on slope (%) | +6.18 (+2.06 – +10.3) |
| Effect of culture-age 11 or 21 days on asymptote (%) | +23.6 (+6.02 – +41.2) |

<sup>a</sup> 95% confidence interval obtained by sampling importance resampling (SIR) using Pearl- Speaks-NONMEM.

## Conclusion

As an *in vitro* culture ages, the proportion of cells that are differentially recovered in liquid media increases as is reflected in the decrease of TTP and TTG for the same CFU recovered on solid media. This indicates that the physiological state alone, in the absence of external stressors, can influence the relative proportions of cells within the parent culture. In the context of disease, it is believed that the bacilli are in a mixed population comprised of cells in different phases of growth. There have been studies indicating that within the population, a majority of cells are either in stationary phase or remain dormant.^9^ This is in line with findings that there is higher recovery of viable bacteria in liquid rather than solid media.^7^

Interestingly, although the presence or absence of drugs did not have any impact on the relationship between TTG and CFU or the GU, drug-treated cultures had a shorter TTP than control for the same CFU. This suggests that TTP identifies a population missed by CFU and is a better estimate for the efficacy of treatment.

To contribute to facilitating extrapolation of our findings towards in-vivo settings, in sputum from patients, future experiments at different concentrations of drugs can confirm the relationship between TTG and CFU/GU is not influenced by exposure gradients.

Here, we demonstrate the existence of bacterial subpopulations that respond differentially to antibiotic treatment, a deeper understanding of the mechanisms involved in switching between subpopulations would assist in the design of antimycobacterial regimens specifically targeted to those that are refractory to treatment and thus leading to poor outcomes.

## Transparency declaration

None to declare.

## Author contributions

FK, PD and TMcH designed and conceptualized the study. HM and AM performed the *in vitro* experiments and collected data. MW performed the modelling analyses. MW and AM wrote the first draft of the manuscript. All authors have read and agreed to the final version of the manuscript.

## References

1. WHO. Global tuberculosis report 2023. 2023.

2. Jindani A, Aber VR, Edwards EA, Mitchison DA. The early bactericidal activity of drugs in patients with pulmonary tuberculosis. Am Rev Respir Dis 1980; 121: 939–49.

3. Mitchison DA, Strum AW. The measurement of early bactericidal activity. Baillières Clin Infect Dis 1997; 4: 185–206.

4. Diacon AH, Maritz JS, Venter A, et al. Time to detection of the growth of Mycobacterium tuberculosis in MGIT 960 for determining the early bactericidal activity of antituberculosis agents. Eur J Clin Microbiol Infect Dis 2010; 29: 1561–5.

5. Diacon AH, Maritz JS, Venter A, Van Helden PD, Dawson R, Donald PR. Time to liquid culture positivity can substitute for colony counting on agar plates in early bactericidal activity studies of antituberculosis agents. Clin Microbiol Infect 2012; 18: 711–7.

6. Bark CM, Okwera A, Joloba ML, et al. Time to detection of Mycobacterium tuberculosis as an alternative to quantitative cultures. Tuberculosis 2011; 91: 257–9.

7. Phillips PPJ, Mendel CM, Nunn AJ, et al. A comparison of liquid and solid culture for determining relapse and durable cure in phase III TB trials for new regimens. BMC Med 2017; 15: 1–9.

8. Dhillon J, Fourie PB, Mitchison DA. Persister populations of Mycobacterium tuberculosis in sputum that grow in liquid but not on solid culture media. J Antimicrob Chemother 2014; 69: 437–40.

9. Mukamolova G V, Turapov O, Malkin J, Woltmann G, Barer MR. Resuscitation-promoting factors reveal an occult population of tubercle bacilli in sputum. Am J Respir Crit Care Med 2010; 181: 174–80.

10. Bowness R, Boeree MJ, Aarnoutse R, et al. The relationship between Mycobacterium tuberculosis MGIT time to positivity and cfu in sputum samples demonstrates changing bacterial phenotypes potentially reflecting the impact of chemotherapy on critical sub-populations. J Antimicrob Chemother 2015; 70: 448–55.

11. Boeckmann AJ, Sheiner LB, Beal SL. NONMEM User’s Guide, Part V. Introductory Guide. 2011: 48.

12. Keizer RJ, Karlsson MO, Hooker A. Modeling and simulation workbench for NONMEM: tutorial on Pirana, PsN, and Xpose. CPT pharmacometrics Syst Pharmacol 2013; 2: 1–9.

13. Lindbom L, Ribbing J, Jonsson EN. Perl-speaks-NONMEM (PsN)—a Perl module for NONMEM related programming. Comput Methods Programs Biomed 2004; 75: 85–94.

14. Chambers JM. Software for data analysis: programming with R. Springer; 2008.

15. Kieser KJ, Rubin EJ. How sisters grow apart: mycobacterial growth and division. Nat Rev Microbiol 2014; 12: 550–62.

16. Aldridge BB, Fernandez-Suarez M, Heller D, et al. Asymmetry and aging of mycobacterial cells lead to variable growth and antibiotic susceptibility. Science (80-) 2012; 335: 100–4.

17. Joyce G, Williams KJ, Robb M, et al. Cell division site placement and asymmetric growth in mycobacteria. 2012.

18. Singh B, Nitharwal RG, Ramesh M, Pettersson BMF, Kirsebom LA, Dasgupta S. Asymmetric growth and division in M ycobacterium spp.: compensatory mechanisms for non-medial septa. Mol Microbiol 2013; 88: 64–76.

19. Santi I, Dhar N, Bousbaine D, Wakamoto Y, McKinney JD. Single-cell dynamics of the chromosome replication and cell division cycles in mycobacteria. Nat Commun 2013; 4: 2470.

20. Avery S V. Microbial cell individuality and the underlying sources of heterogeneity. Nat Rev Microbiol 2006; 4: 577–87.

21. Dhar N, McKinney J, Manina G. Phenotypic heterogeneity in Mycobacterium tuberculosis. Tuberc Tuber Bacillus 2017: 671–97.

