## Supplemental data for "The impact of physiological state and environmental stress on bacterial load estimation methodologies for *Mycobacterium tuberculosis*"

**Description of the models used in the study**

A. **Analysis of relationship between TTP and CFU**

A linear model estimating an intercept and slope adequately described the relationship between TTP and CFU (Figure 1). Typical values for intercept and slope were estimated to 215 hours and -50.3 hours/log10 CFU, respectively (Table 1). Between-biological replicate variability was included on the intercept and an additive error model on logarithmic scale described the residual variability. Additive error was inflated by 1.94-fold for samples with <10 colonies (dOFV = -22.8, degrees of freedom, df = 1, p < 0.001). The presence of drug significantly affected the intercept, with drug exposure resulting in 17.6% lower intercept (dOFV = -7.33, df = 1, p = 0.007). Samples cultured on day 11 or 21 had 15.3% lower intercept (dOFV = -11.3, df = 1, p < 0.001) compared to samples cultured on day 0 or 3. A VPC stratified by drug presence and culture-age can be found in Figure 1. Similar results were found in the dataset with imputed CFUs (Figure S1, Table S1), with the difference of samples with culture-age of 0, 11 or 21 days had 18.6-folds larger random variability than samples with culture-age of 3 days (difference in OFV, dOFV = -7.92) in the data with no imputations.

B. **Analysis of relationship between TTG and CFU**

The relationship between TTG and CFU was adequately described by a linear model estimating an intercept and slope to 184 hours and -42.2 hours/log10 CFU, respectively (Figure 2 and Table 2). Between-biological replicate variability was included on the intercept and an additive error model on logarithmic scale was used to describe the residual variability. Culture-age significantly affected TTG for the same CFU with 18.3% lower intercept for culture ages of 11 and 21 days than that for those 0 and 3 days (dOFV = -25.7, df = 1, p < 0.001). The presence of drugs did not have any impact on the relationship between TTG and CFU. A VPC stratified by culture-age can be found in Figure 2. Similar results were found in the dataset with imputed CFUs (Figure S2, Table S2).

C. **Analysis of relationship between GU and time**

A total of 184,129 observations from 227 experiments were included in the analysis (Figure 3). A logistic model estimating an asymptote, a slope, and the time of the function’s midpoint (T50) fit the data best. Typical values for the asymptote, slope and T50 were 12,800 GU, 0.069 GU/hour and 88.9 hours, respectively (Table 3). Random variability was included on all three parameters and an additive error model on logarithmic scale described the residual variability. Estimating covariance between the random variabilities in the slope and T50 further improved the model fit (dOFV = -445). Culture-age significantly affected the slope and asymptote, with samples cultured on day 11 or 21 having 6.18% bigger slope (dOFV = -8.15, df = 1, p = 0.004) and 23.6% higher asymptote (dOFV = -8.78, df = 1, p = 0.003) compared to samples cultured on day 0 or 3. A VPC stratified by culture-age is shown in Figure 3.

**Figures and Tables**

Table S1: Parameter estimates for model describing relationship between TTP and CFU in dataset with imputations.

| **Parameter** | **Typical value (95% CI)^a^** |
| --- | --- |
| Intercept (hours) | 228 (209 – 250) |
| Slope (hours/log10 CFU) | -28.7 (-31.3 – -26.2) |
| Additive error on log scale (%) | 13.9 (12.5 – 15.7) |
| Scaling of additive error on log scale for < 10 CFU (folds) | 1.90 (1.54 – 2.43) |
| Between-biological replicate variability on intercept (%CV) | 9.54 (6.52 – 12.4) |
| Effect of drug presence on intercept (%) | -21.7 (-34.5 – -9.02) |
| Effect of culture-age 11 or 21 days on intercept (%) | -10.4 (-13.9 – -6.87) |

^a^ 95% confidence interval obtained by sampling importance resampling (SIR) using Pearl-Speaks-NONMEM.

Table S2: Parameter estimates for model describing relationship between TTG and CFU in dataset with imputations.

| **Parameter** | **Typical value (95% CI)^a^** |
| --- | --- |
| Intercept (hours) | 186 (172 – 200) |
| Slope (hours/log10 CFU) | -30.9 (-34.3 – -27.2) |
| Additive error on log scale (%) | 20.2 (18.3 – 22.8) |
| Between-biological replicate variability on intercept (%CV) | 7.07 (4.34 – 9.50) |
| Effect of culture-age 11 and 21 days on the intercept (%) | -13.0 (-18.3 – -7.68) |

^a^ 95% confidence interval obtained by sampling importance resampling (SIR) using Pearl-Speaks-NONMEM.

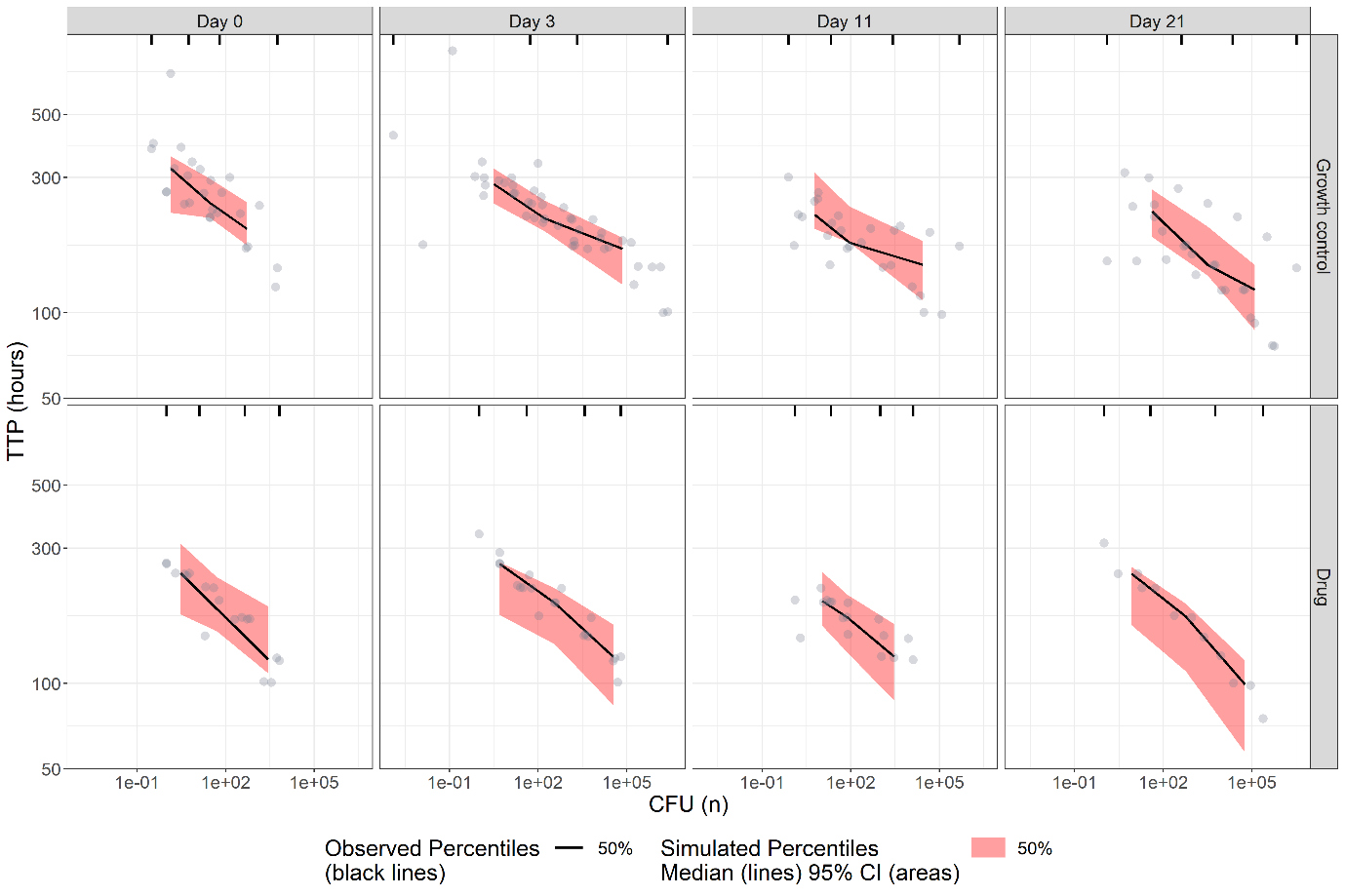

Figure S1. Visual predictive check (n=1000) of TTP versus CFU in dataset with imputations, stratified by culture-age and presence of drug. The circles represent observations.

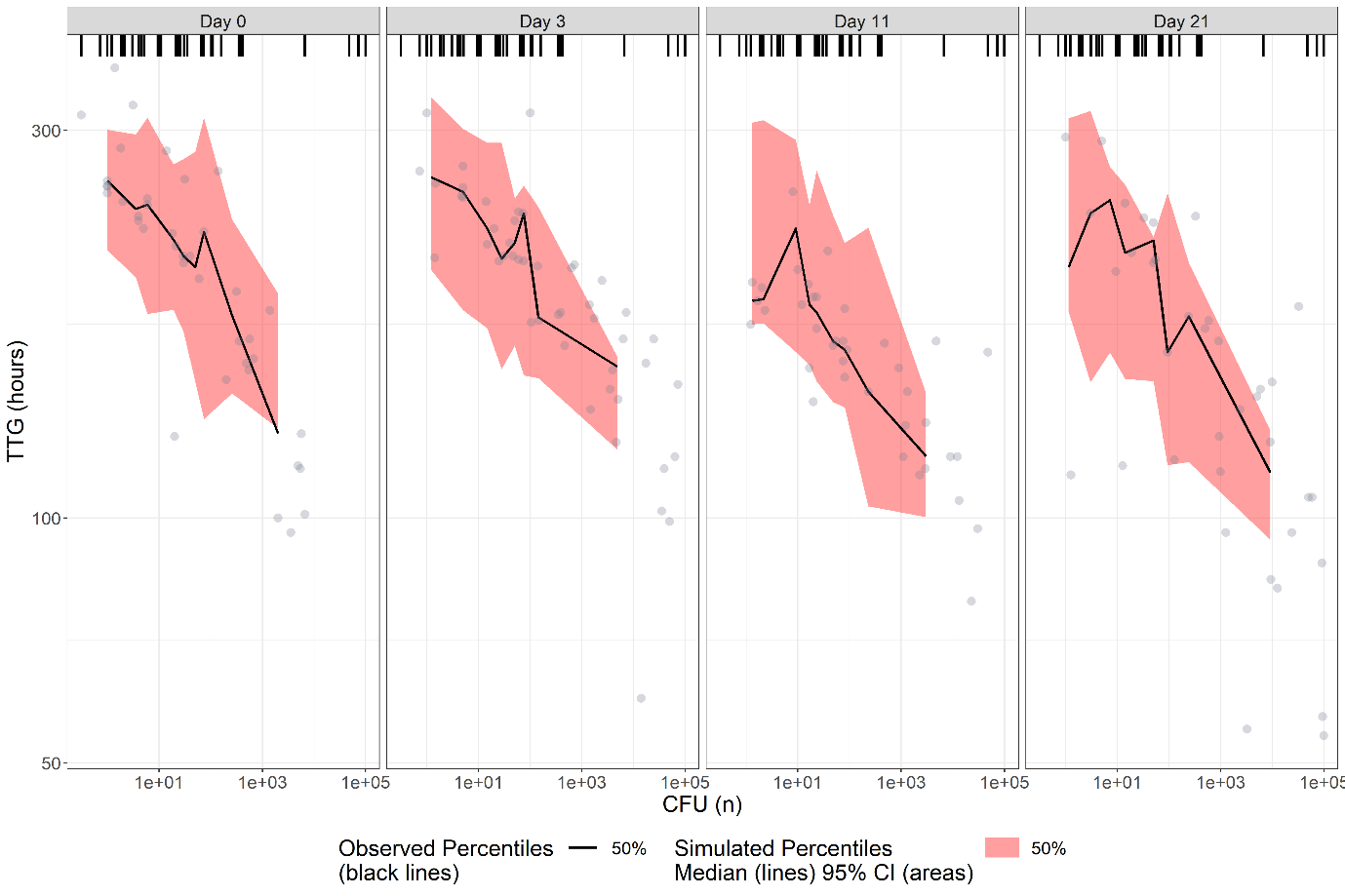

Figure S2. Visual predictive check (n=1000) of TTG versus CFU in dataset with imputations, stratified by culture-age. The circles represent observations.

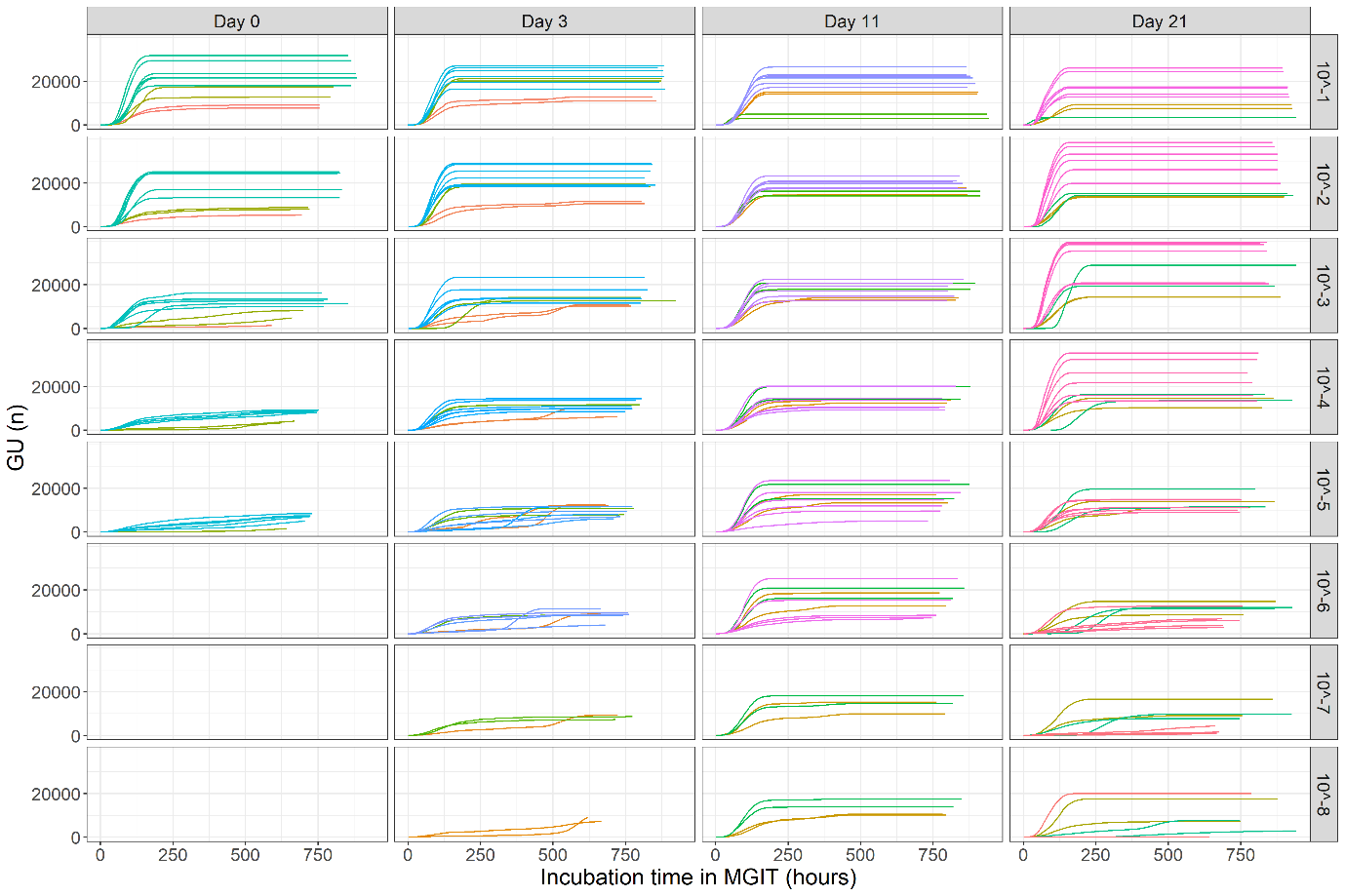

Figure S3. Raw GU data stratified by culture-age and dilution. Each trajectory represents GU reads from one MGIT.

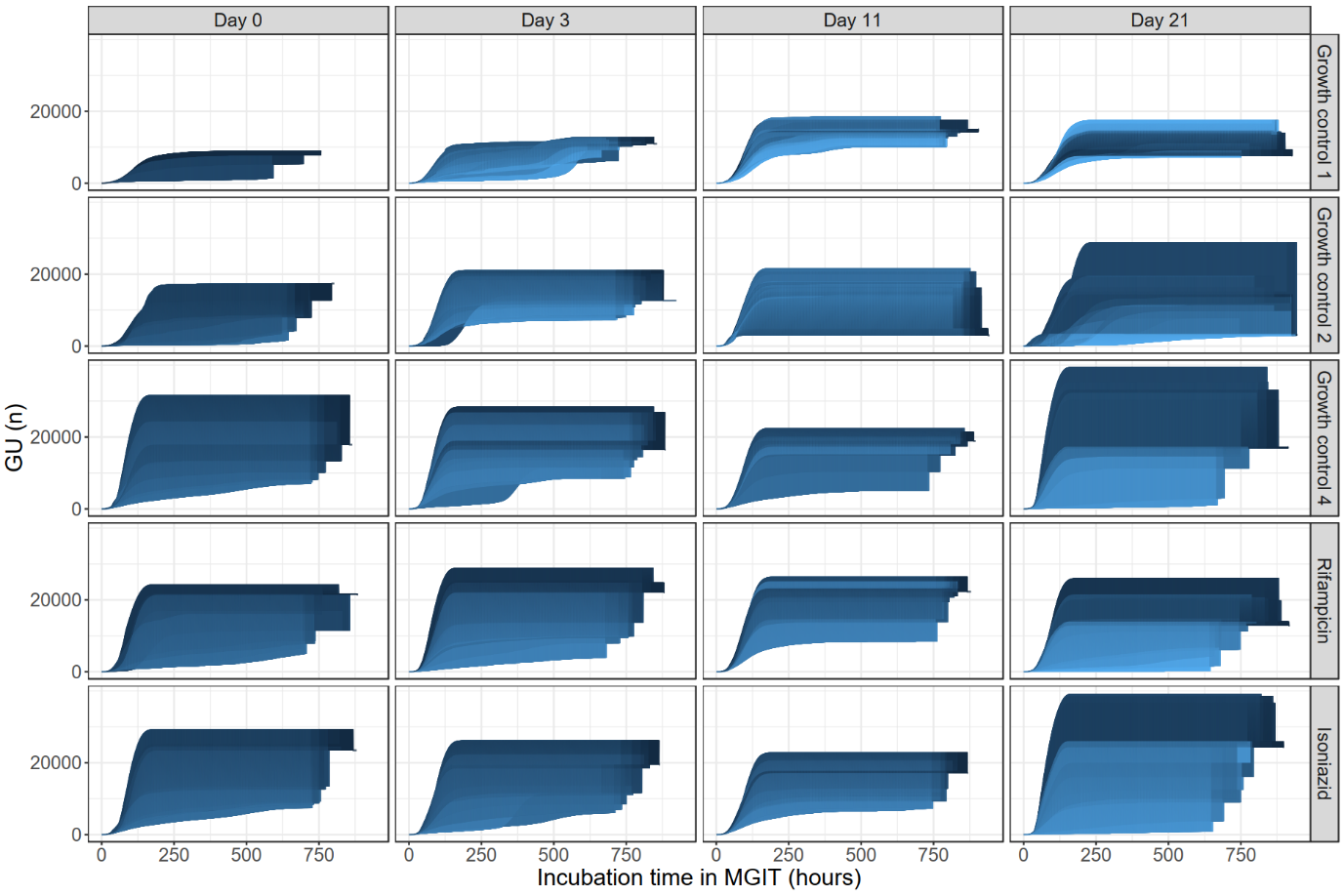

Figure S4. Raw GU data stratified by culture-age and biological replicate and coloured by dilution, where the lighter lines represent more diluted samples. Each trajectory represents GU reads from one MGIT.
